# A new *Oprm1*-Cre mouse line enables detailed molecular characterization of μ-opioid receptor cell types

**DOI:** 10.1101/2022.06.09.495430

**Authors:** Juliet Mengaziol, Amelia D. Dunn, Jordi Crues-Muncunill, Darrell Eacret, Chongguang Chen, Kathryn Bland, Lee-Yuan Liu-Chen, Michelle E. Ehrlich, Julie A. Blendy

## Abstract

Key targets of both the therapeutic and abused properties of opioids are μ-opioid receptors (MORs). Despite years of research investigating the biochemistry and signal transduction pathways associated with MOR activation, we do not fully understand the cellular mechanisms underlying opioid addiction. Given that addictive opioids such as morphine, oxycodone, heroin, and fentanyl all activate MORs, and current therapies such as naloxone and buprenorphine block this activation, the availability of tools to mechanistically investigate opioid-mediated cellular and behavioral phenotypes are necessary. Therefore, we derived, validated, and applied a novel MOR-specific Cre mouse line, inserting the Cre coding sequence into the MOR 3’UTR to generate a bicistronic gene. As intended, there were no differences in behavioral responses to morphine when compared to wild type mice, nor are there any alterations in *Oprm1* gene expression or receptor density. To assess Cre recombinase activity, MOR-Cre mice were crossed with the floxed GFP-reporters, Rosa^LSLSun1-sfGFP^ or Rosa^LSL-GFP-L10a^. The latter allowed for cell type specific RNA sequencing via TRAP (Translating Ribosome Affinity Purification) of striatal MOR+ neurons following opioid withdrawal. This new tool will facilitate the study of opioid biology under varying conditions.

## Introduction

In the past 20 years, the opioid use and overdose crisis in the US has ignited a resurgence of research in opioid biology. Both endogenous and exogenous (including illicit) opioids such as morphine, heroin, oxycodone, and fentanyl exert their effects primarily through the G-protein coupled μ-opioid receptor (MOR) which is encoded by the *OPRM1* gene. MORs are necessary for the effects of opioid induced analgesia, reward, dependence and opioid withdrawal [1–3]. Early autoradiography studies have demonstrated that MORs are distributed across the central nervous system, with a high concentration in brain regions known to be involved in sensorimotor integration, pain and reward such as the caudate-putamen, thalamus, dorsal horn of the spinal cord, ventral tegmental area and nucleus accumbens [4–8]. The fact that the MOR is critical for the analgesic and addictive properties of opioids is well established. However, the MOR-expressing neuronal populations and circuits that mediate these effects have not been fully or precisely mapped. Moreover, while the signaling pathways downstream of opioid binding to MORs have been delineated *in vitro,* the *in vivo* functions have not been studied specifically in *Oprm1*-expressing neurons. Therefore, large gaps exist in our understanding of the molecular processes initiated by opioids in a cell-type specific fashion.

We describe herein the derivation and characterization of a mouse that expresses a T2A-cleaved Cre-recombinase under the control of the *Oprm1* gene promoter and enhancers. We show that insertion of the T2A-Cre gene into the 3’UTR of the *Oprm1* locus does not disrupt normal *Oprm1* gene expression or MOR levels in the brain. Additionally, we demonstrate that *Oprm1*-Cre mice (Oprm1^Cre/Cre^ or Oprm1^Cre/+^) respond identically to wild-type mice (Oprm1^+/+^) in behavioral measurements of both acute and chronic morphine administration. We show how this mouse can be used in conjunction with additional genetic lines for the visualization and molecular characterization of *Oprm1*-expressing cells and employ this new tool to delineate the transcriptional changes in *Oprm1*-expressing striatal cells during spontaneous morphine withdrawal. Overall, this study demonstrates the reliability and utility of this *Oprm1*-Cre for the detailed study of *Oprm1*-expressing cells in the mouse brain.

## Materials and methods

### Drugs

Morphine sulfate was obtained from the NIDA Drug Supply (Research Triangle Park, NC) and dissolved in 0.9% saline. Naloxone hydrochloride dyhydrate was purchased from Sigma.

#### Animals

C57Bl/6NTac male and female mice were mated for embryo recovery and injections. MOR-Cre mice were crossed with either *Rosa26^LSL-GFP-L10a^* or *Rosa^LSLSun1-sfGFP^* [9] and maintained on a C57BL/6NTac background. These reporter mice were used to confirm the functionality of Cre-mediated recombination and eventually for TRAP and RNA sequencing experiments. All behavior and molecular studies were conducted in male mice which were group-housed with food and water available *ad libitum* and maintained on a 12 h light/dark cycle, lights on at 6:00am, according to the University of Pennsylvania Animal Care and Use Committee. Mice weighed 20–30 g and were 8–14 weeks old.

### Genome editing procedure

We employed CRISPR-Cas9 assisted gene targeting in mouse C57Bl/6 embryos for the rapid derivation of the *Oprm1*-Cre line. A targeting construct encoding a T2A cleavable peptide and Cre recombinase was inserted downstream of the last exon (exon 4) in the *Oprm1* gene sequence. Genome-editing using CRISPR-Cas9 technology has been shown to have the propensity for off-target effects, most of which occur at sequence elements with high similarity to the target, containing at most two mismatches to the guide RNA. Therefore, we utilized the “CRISPR design” algorithm, developed by the Zhang lab (http://crispr.mit.edu), to select the optimal pair of single-stranded guide RNAs (sgRNAs) to target the *Oprm1* 3’ UTR. We then combined the Cas9 ‘nickase’, the D10A mutant version of the protein which can only cleave one of the two DNA strands, with a pair of offset guide RNAs directed to opposite strands of the target sequence. Lastly, the inhibitor of non-homologous end-joining, SCR7, was used to increase the frequency of homology-based repair using the provided donor template at the expense of NHEJ [10].

The two guide RNAs were synthesized by *in vitro* transcription using T7 RNA polymerase (sequence available upon request). The T7 promoter and sgRNA sequences were amplified by PCR, resulting in a ~ 120 bp template, which was gel-purified before use in the *in vitro* transcription reaction. The sgRNAs were purified using an RNA affinity filter (MEGAclear) and quantified using capillary electrophoresis (Bioanalyzer). This also served as a quality control step to ascertain that the sgRNAs synthesized were full length. One hour before the microinjection procedure, an ‘injection mix’ was prepared containing the two sgRNAs at a final concentration of 50 ng/μl, biotinylated targeting template 20 ng/μl and mRNA encoding the Cas9-mSA nickase at 75 ng/μl. The injection mixture was microinjected into C57BL/6NTac two-cell embryos and cultured *in vitro* in the presence of 50 nM SCR7. Embryos were placed in this environment for 24 hours to allow development to the morula stage (8 to 16 cell embryo) in order to inhibit NHEJ and increase the frequency of homology directed repair (HDR) [10]. Embryos were then implanted into the uterus of pseudo pregnant mice and potential founder mice born three weeks later. To account for off-target mutagenesis, the founder lines established were analyzed in relation to C57Bl/6NTac for any spurious changes in other parts of the genome by selecting the top five most similar off-target sites identified by “CRISPR design,” and assessed their potential mutations by Sanger sequencing of the relevant PCR amplicons. For the *Oprm1*-Cre line, one founder line was selected that passed this test and genome sequencing was performed at 50x coverage to identify any mutations relative to the C57BL/6 reference genome. We found no off-target functional variants in the targeted mice.

### PCR and primers for genotyping of novel mouse line

*Primers:* Forward primer for the *Oprm1*-T2ACre (Oprm1 ^Cre/+^ or Oprm1^Cre/Cre^) allele: 5’ CGCTGGAGTTTCAATACC GG 3’; Forward primer for wild type (Oprm1^+/+^) allele: 5’ ACTGCTCCATTGCCCTAACT 3’; Common Reverse Primer: 5’ TGACGTCCGGTGATGACTTA 3’. *Thermocycler conditions:* 95°C for 5 min; 40 cycles of 1. 95°C for 45 sec, 2. 60°C for 45 sec, 3. 72°C for 1:30 min; 72°C for 10 min; 4°C infinite. *Expected PCR products:* 273 bp (1 band, homozygous for wild type), 350bp (1 band, homozygous for *Oprm1*-T2ACre knock in), 237bp & 350bp (2 bands, heterozygous).

### Brain dissection and collection

Mice were cervically dislocated and brains were quickly extracted, and dissected on ice (hypothalamus, cerebellum, hippocampus, striatum, prefrontal cortex, cortex, and thalamus). Tissue was collected in microtubes, flash frozen in liquid nitrogen and stored in a –80°C.

### Receptor binding

Five hundred mL of the lysis buffer (50 mM Tris-HCl buffer, 0.1 mM phenylmethylsulfonyl fluoride, pH 7.4) was added to one tissue sample in a tube and sonicated 5 sec on ice at low power. Protein concentration was determined by the BCA method and adjusted to 150 mg/0.1ml. Binding was performed with the selective MOR agonist [^3^H]DAMGO (53.7 Ci/mmole, Perkin Elmer, Boston, MA) in 50 mM Tris-HCl buffer (pH 7.4). Each binding assay was carried out in duplicate in a final volume of 1 ml with 1.65 nM [^3^H]DAMGO +/− naloxone (10 mM) and 150 mg protein. After incubation for 1 hour at room temperature, bound and free ligand were separated by rapid filtration over GF/B filters with a 24-sample cell harvester (Brandel, Gaithersburg, MD) under vacuum. Filters were washed four times with ice-cold 50 mM Tris-HCl buffer. Radioactivity trapped in the filters was determined by liquid scintillation counting with a b-counter. The counting efficiency was approximately55%.

### RNA isolation and *Oprm1* expression

RNA was isolated from each brain region using the Absolutely RNA Miniprep Kit (Agilent Technologies, Cat. No. 400800), and stored at —80°C [11]. mRNA was diluted such that 200ng was used to synthesize cDNA using the High-Capacity cDNA Reverse Transcription Kit (Applied Biosystems, Cat. No. 4368814). The reverse-transcription recycling parameters were set as: 25°C for 10 min, 37°C for 2 hours, 85°C for 5 min, and 4°C until next step. For quantitative real-time polymerase chain reactions (qPCR), SYBR Green qPCR reactions were assembled using SYBR™ Green PCR Master Mix (Applied Biosystems, Cat. No. 4309155) along with the final concentration of 300 nM for each of the primers. All qPCR reactions were run using the Stratagene MX3000 and MXPro QPCR software under the following cycle conditions: 95°C for 10 min, followed by 40 cycles of 95°C for 15 sec and 60°C for 1 min. All reactions were performed in triplicate and the average cycle threshold was used for analysis. The mRNA levels of Oprm1 were normalized to the ‘housekeeping’ gene, Glyceraldehyde-3-Phosphate Dehydrogenase (GAPDH). *Primers:* Spanning Exons 3 and 4 of Oprm1: 5’ TCCCAACTTCCTCCACAATC; 3’ TAGGGCAATGGAGCAGTTTC. GAPDH: 5’ AACGACCCCTTCATTGACCT; 3’ TGGAAGATGGTGATGGGCTT (Eurofins).

### Acute morphine-induced locomotor response

Locomotor activity of both Oprm1^Cre/+^ and Oprm1^+/+^ animals was analyzed using the Continuous Open Mouse Phenotyping of Activity and Sleep Status (COMPASS) system [12]. Mice were removed from their grouped home cage and singly housed in clean home cages (26 x 20 x14 cm) without nestlets. Data analysis using pyroelectric or passive infrared sensors (PIRs) attached to the tops of the testing cages and began once mice were placed into testing cages for 30 minutes of habituation. After habituation, each mouse was briefly removed from the home cage and a single 20 mg/kg i.p. injection of morphine was administered to measure acute locomotor response. Individual activity was recorded for 2 hours post morphine injection, totaling a testing time of about 2 hours and 40 minutes per mouse.

### Cumulative hot plate analgesia

*Oprm1*^Cre/+^ *and Oprm1*^+/+^ animals were injected with increasing doses of morphine (0, 1, 2, 7, 20 mg/kg, i.p.) in 30-minute intervals. After each injection, mice were placed on a 55°C hot plate and latency to lick the hind paw and jump were recorded. Upon displaying one or both of these behaviors before the predetermined cut off time (60 sec), mice were removed from the hot plate until the next injection.

### Chronic morphine-induced dependence & precipitated withdrawal

Both *Oprm1*^Cre/+^ and *Oprm1^+/+^* animals underwent a chronic-morphine exposure paradigm to induce dependence using repeated, subcutaneous (s.c.) injections of escalating doses of morphine (20mg/kg; 40mgkg; 60mg/kg; 100mg/kg, injections, twice /day injections for 5 days) [13]. Two hours after the last 100mg/kg injection on day 5 of the exposure paradigm, animals were placed on cotton pads inside of an open-topped clear plastic cylinder and a single dose of 1mg/kg of naloxone was administered to each animal after a 30-minute habituation period. Somatic signs of withdrawal were scored in 5-minute bins for 30 minutes with a measurement of the following behaviors: Ptosis (1/min), resting tremor (1/min), diarrhea (1/min), teeth chatter (1/min), genital licking, gnawing, head & body shakes, paw tremors, scratches, backing, and jumping. A cumulative withdrawal score per animal was calculated by tallying all instances of somatic signs in the 30-minute test [11]. For the Oprm1^Cre/Cre^ group that underwent spontaneous withdrawal for TRAP RNA sequencing analysis, animals did not receive any naloxone and brains were collected 24 hours after (on day 6) the last 100mg/kg injection.

### Tissue processing and image acquisition

*Oprm1*^Cre/Cre^; Rosa^LSL-GFP-L10a^ and *Oprm1*^Cre/Cre^; Rosa^LSLSun1-sfGFP^ mice were anesthetized and intracardiac perfusions performed by hand (10mL/min) with 35mL of chilled 1x PBS (Roche) followed by 50mL of freshly made, chilled 4% paraformaldehyde (PFA) (Sigma-Aldrich). After collection, whole brains were post fixed for 12-24 hours in 4% PFA solution at 4°C. Brains were then placed in fresh, chilled 30% sucrose at 4°C until tissue density equilibrated (2-3 days). Brains were rapidly frozen at-80°C and 30-micron sections were cut on a cryostat at 20°C (Cryostar NX70). Sections were mounted (Fisherbrand superfrost slides) with DAPI Fluoromount slide cover sealer (SouthernBiotech). Images were acquired as 1-μm z-stacks at 20× magnification on a Keyence BZ-X800 fluorescence microscope and stitched to capture entire brain regions within the coronal plane. Regions of interest were defined according to the Allen Mouse Brain Atlas.

### Translating ribosomal affinity purification (TRAP) assay

RNA was isolated from striatal cells expressing GFP-L10a by TRAP [9]. Briefly, 24 hours after the last chronic morphine (100mg/kg) or saline dose, Oprm1^Cre/Cre^; Rosa^LSL-GFP-L10a^ mice were euthanized via cervical dislocation, and striatum was isolated (both hemispheres, 2 brains pooled per sample), homogenized with lysis buffer, and incubated with magnetic beads that were pre-conjugated to anti-GFP antibodies (clones htz-GFP-19F and htz-GFP-19C8, Memorial Sloan-Kettering Monoclonal Antibody Facility, New York, New York, USA) to affinity purify RNA that was bound by GFP-L10a fusion protein[14].

### RNA-sequencing and analysis

RNA integrity was assessed with an Agilent Tapestation 4200 using the RNA ScreenTape (Agilent, Wilmington, DE). All samples had RIN values > 8. mRNA libraries were generated using the NEBNext Ultra™ II RNA Library Prep Kit for Illumina (New England BioLabs, Ipswitch, MA). Resulting libraries were sequenced on a NovaSeq 6000 at a depth of 30 million reads/sample, with paired-end sequencing of 150 base pairs (PE150). Mapping of the reads to the mouse reference genome *(Mus musculus,* GRCm38/mm10) was performed using the STAR aligner (v2.6.1d). Differential expression analysis was performed with DESeq2 (v1.20.0). Differential expression cut-offs were set at an adjusted (Adj.) *p*-value < 0.05 and a Log2 fold change (Log2FC) of 0.5. Gene ontology (GO) analyses were performed via goProfiler, and the significant GO terms (p<0.05) were reduced using Revigo. Enrichment for cell-type specific gene sets was assessed using a hypergeometric test.

## Results

### The T2ACre sequence insertion into the Oprm1 locus does not alter Oprm1 expression or MOR levels

CRIPSR/Cas9 mediated homologous recombination allows for precise gene targeting of *Oprm1*. Sequencing analysis confirmed the appropriate T2A-Cre-recombinase insertion into the *Oprm1* gene locus (Fig 1B). Genotyping primers were designed to distinguish wild type and Cre-inserted alleles (Fig 1C). Quantitative-PCR (qPCR) results confirm that *Oprm1*^Cre/+^ and *Oprm1*^+/+^ (wild type, or WT) animals have comparable Oprm1 transcript levels across several brain regions (Fig 2A). There was no difference in *Oprm1* mRNA levels (2-way ANOVA revealed a nominal effect of genotype between groups (F(1,60) = 0.5438, p = 0.4637)) in the hippocampus, hypothalamus, prefrontal cortex, cerebellum, striatum, or thalamus. At the protein level, specific [^3^H]DAMGO binding to the MOR was not significantly altered in the cortex, hippocampus, or thalamus (Fig 2B). Two-way ANOVA revealed a main interaction effect [F(2, 32) = 3.306, p =0.0495], but post hoc test revealed no difference between genotypes in any brain region.

**Fig 1.**
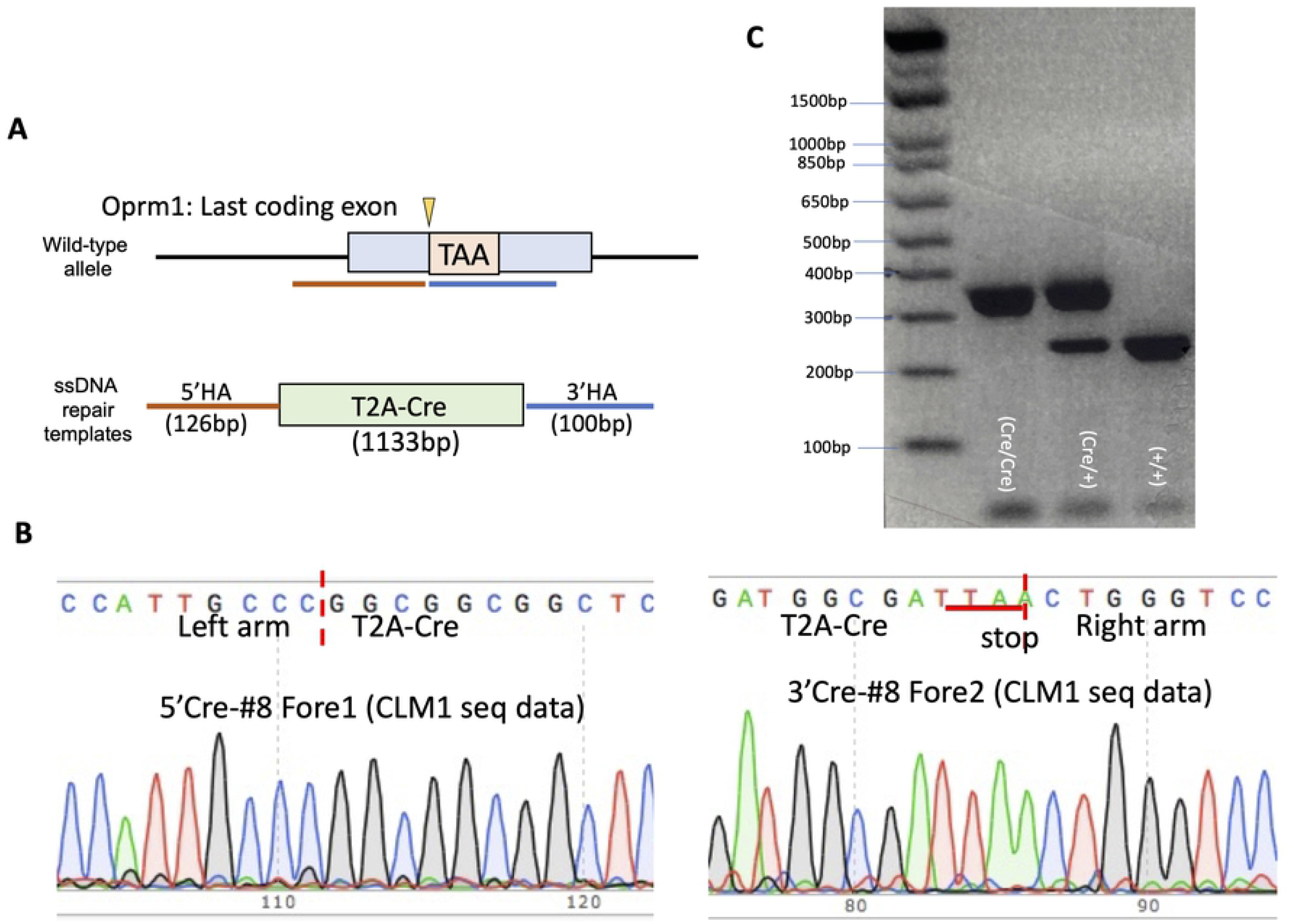
Genetic construct generation and sequencing of the *Oprm1-Cre* mouse line. (A) Cartoon representation of the genetic targeting strategy to create the desired mouse line. CRISPR/Cas9 mediated insertion encoding a functional Cre-recombinase (Cre) enzyme was inserted upstream of the Oprm1 stop codon, in Exon 4. A T2A-cleavable peptide was included in this genetic construct to allow for Cre release from the mu-opioid receptor (MOR) once the entire Oprm1-T2ACre gene is translated to avoid unwanted protein interactions. (B) Confirmational DNA sequencing of the T2ACre genetic insert. (C) Genotyping PCR products for homozygote (Oprm1^Cre/Cre^) heterozygote (Oprm1^Cre/+^), and wild-type (Oprm1^+/+^) mice. Wildtype allele is 273bp and T2ACre allele is 350bp.

**Fig 2.**
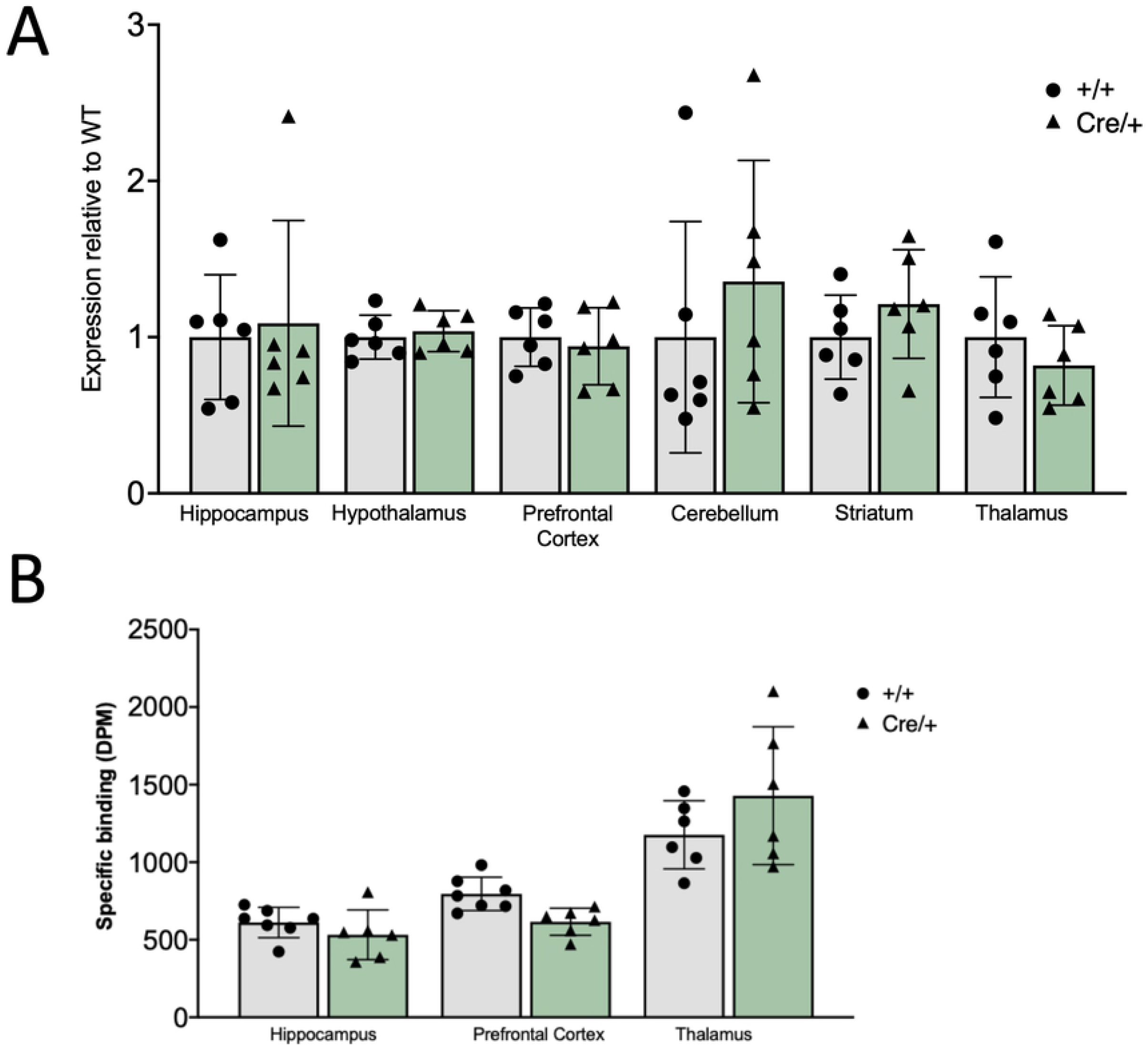
Oprm1 mRNA and 3HDAMGO binding is not altered in the *Oprm1-Cre* mouse line compared to wild type C57Bl/6 mice. Animals expressing the T2A-Cre allele showed no significant molecular differences of Oprm1-expression when compared to wild-type littermates. (A) Quantification of the qPCR analysis of Oprm1 between Oprm1^Cre/+^ and Oprm1^+/+^ animals shows that there is comparable Oprm1 mRNA transcript levels in the hippocampus, hypothalamus, PFC, cerebellum, striatum, and thalamus, regardless of genotype (Males, 10-15wks old, n=6). Two-way ANOVA revealed no main effect of genotype between groups (F (1,60) = 0.5438, p = 0.4637). (B) 3H-DAMGO binding is not different between groups. (Males, 10-15wks old, n=6-7). Two-way ANOVA revealed a main interaction effect (F(2, 32) = 3.306, p = 0.0495. Post hoc test revealed no difference between Oprm1^Cre/+^ and Oprm1^+/+^ mouse lines in any brain region.

### T2ACre sequence insertion into the Oprm1 locus does not alter MOR behavioral function

The effects of both acute and chronic morphine were tested to assess MOR function following insertion of the T2ACre sequence. As expected, locomotor activity increased following an acute morphine (20mg/kg) injection in both *Oprm* ^Cre/+^ and WT animals (Fig 3A). 2-way ANOVA revealed a significant effect of time post-injection on locomotor activity (F_(9, 120)_ = 2.53, p < 0.05), with no effect of genotype (F_(1, 120)_ = 0.60, p = 0.44). Thus, mice with the Cre insert have similar locomotor responses to morphine as the WT mice, with intact acute morphine processing, reactivity, and MOR-function.

**Fig 3.**
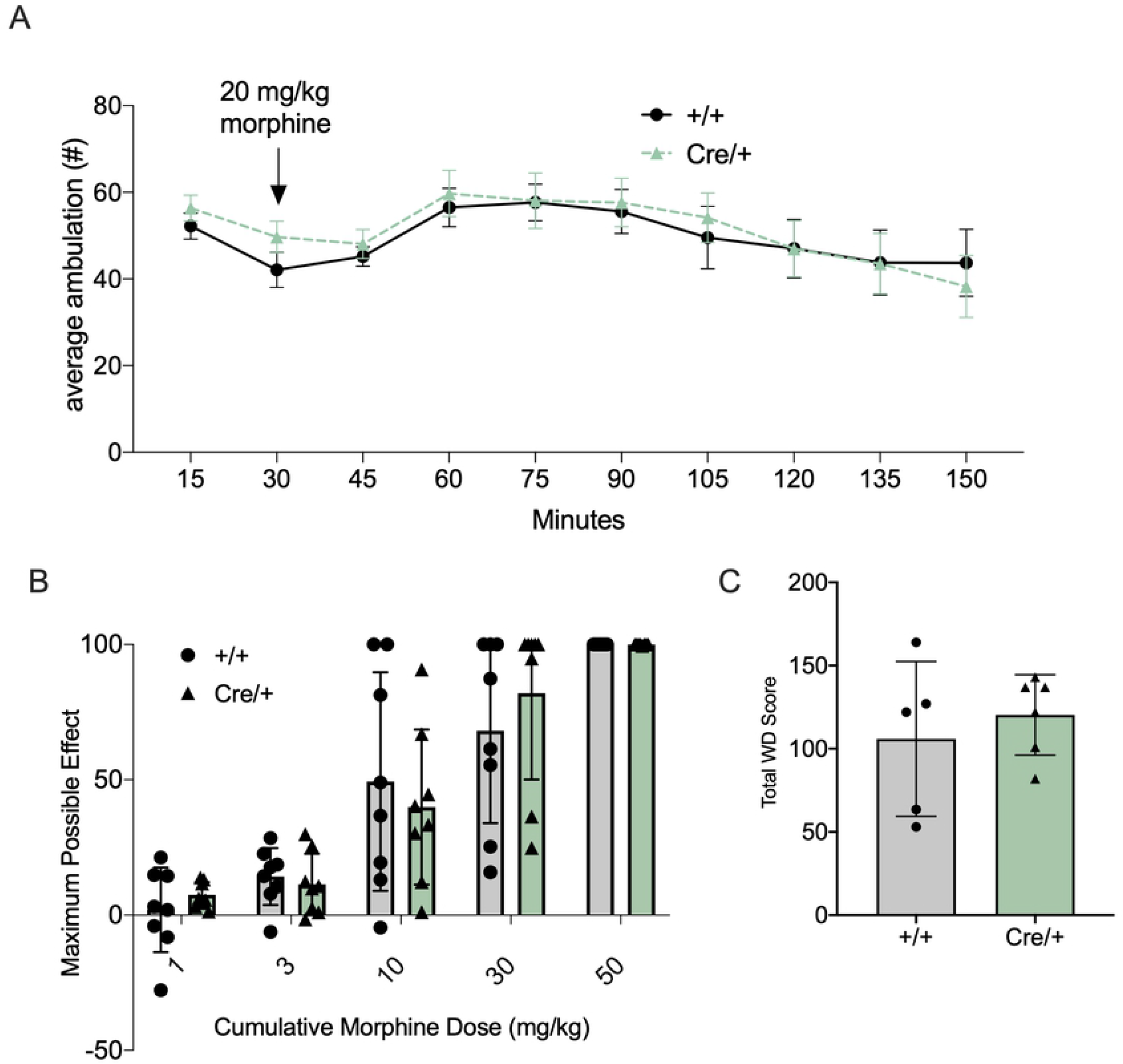
Confirmation of MOR function and acute and chronic opiate-mediated behaviors/phenotypes in the *Oprm1-Cre* mouse line. Animals expressing the T2A-Cre allele showed no significant behavioral differences of morphine-induced phenotypes when compared to wild-type littermates. (A) No significant differences of acute morphine-induced locomotion responses occurred between Oprm1^Cre/+^ and Oprm1^+/+^ animals. Following 20mg/kg morphine injections activity was recorded for 2hrs (males, 10-15wks old, n=8/group) (2-way ANOVA: significant effect of time post-injection on locomotor activity (F_(9, 120)_ = 2.53, p < 0.05), no effect of genotype (F_(1, 120)_ = 0.60, p = 0.44). (B) Morphine-mediated antinociception, as measured by hind-paw lick latency on a 55 °C hot-plate assay using a cumulative-dosing paradigm, was not different (males, 10-15wks old, n=8/group) (2-way ANOVA analysis: significant effect of morphine dose (F_(1.43, 12.89)_ = 6.36, p = 0.0178) and no genotype effect (F_(1,9)_ = 0.016, p =0.901). Results are presented as percentage of maximal possible effect (MPE) [(morph jump latency - saline jump latency)/(total time - saline jump latency) × 100] (mean ± SEM, *n* = 18). (C) Naloxone induced precipitated withdrawal after a chronic morphine exposure paradigm is intact between Oprm1^Cre/+^ and Oprm1^+/+^ animals. Cumulative withdrawal score per animal was calculated by tallying all instances of somatic signs in the 30-minute test interval (males, 10-15wks old, n=5-6) (unpaired two tailed t-test: p = 0.523).

The analgesic effect of morphine was tested using a cumulative morphine hot plate analgesia assay, with a dose range of 1mg/kg morphine to 50mg/kg morphine (Fig 3B). There were no significant differences in the latency to respond between *Oprm1*^Cre/+^ and *Oprm1*^+/+^ animals. A 2-way ANOVA analysis revealed a significant effect of morphine dose (F_(1.43, 12.89)_ = 6.36, p = 0.0178) and no effect of genotype (F_(1,9)_ = 0.016, p =0.901).

To confirm that the effects of chronic opioid exposure were also maintained in *Oprm1*^Cre/+^ mice, dependence, induced following 5 days of morphine exposure, and subsequent withdrawal behavior was measured. 2 hours after the last 100mg/kg morphine injection, animals received 1mg/kg of naloxone i.p. and were observed for somatic withdrawal behaviors. There were no significant differences in behavior between the *Oprm1*^Cre/+^ and *Oprm1*^+/+^ animals (unpaired two tailed t-test: p = 0.523), indicating intact MOR function during morphine dependence and withdrawal (Fig 3C). Together, these behavioral and molecular results indicate that the T2ACre sequence insertion into the Oprm1 locus did not alter MOR expression or function.

### The T2ACre sequence insertion into the Oprm1 locus produces functional Cre-recombinase enzyme in MOR expressing cell-types

We used two floxed GFP-reporters, Rosa^LSL-GFP-L10a^ and Rosa^LSLSun1-sfGFP^ [9], to confirm activity of Cre-mediated recombination. When crossed with the Rosa^LSL-GFP-L10a^ line, recombinant *Oprm1*^Cre/+^; *Rosa*^LSL-GFP-L10a^ mice produce GFP-tagged ribosomal protein L10a in *Oprm1*-expressing cells. At 2X, 10X and 20X imaging, we observed dense regions of GFP in the ventral tegmental area (VTA), the dentate gyrus of the hippocampus, and the periaqueductal gray (PAG), all regions that have been shown to have high MOR expression density in the mouse brain [15–17] (Fig 4). To confim ribosomal-GFP expression, 40X images display that GFP is expressed in the cytoplasm throughout the cell, consistent with GFP-tagged ribosomes.

**Fig 4.**
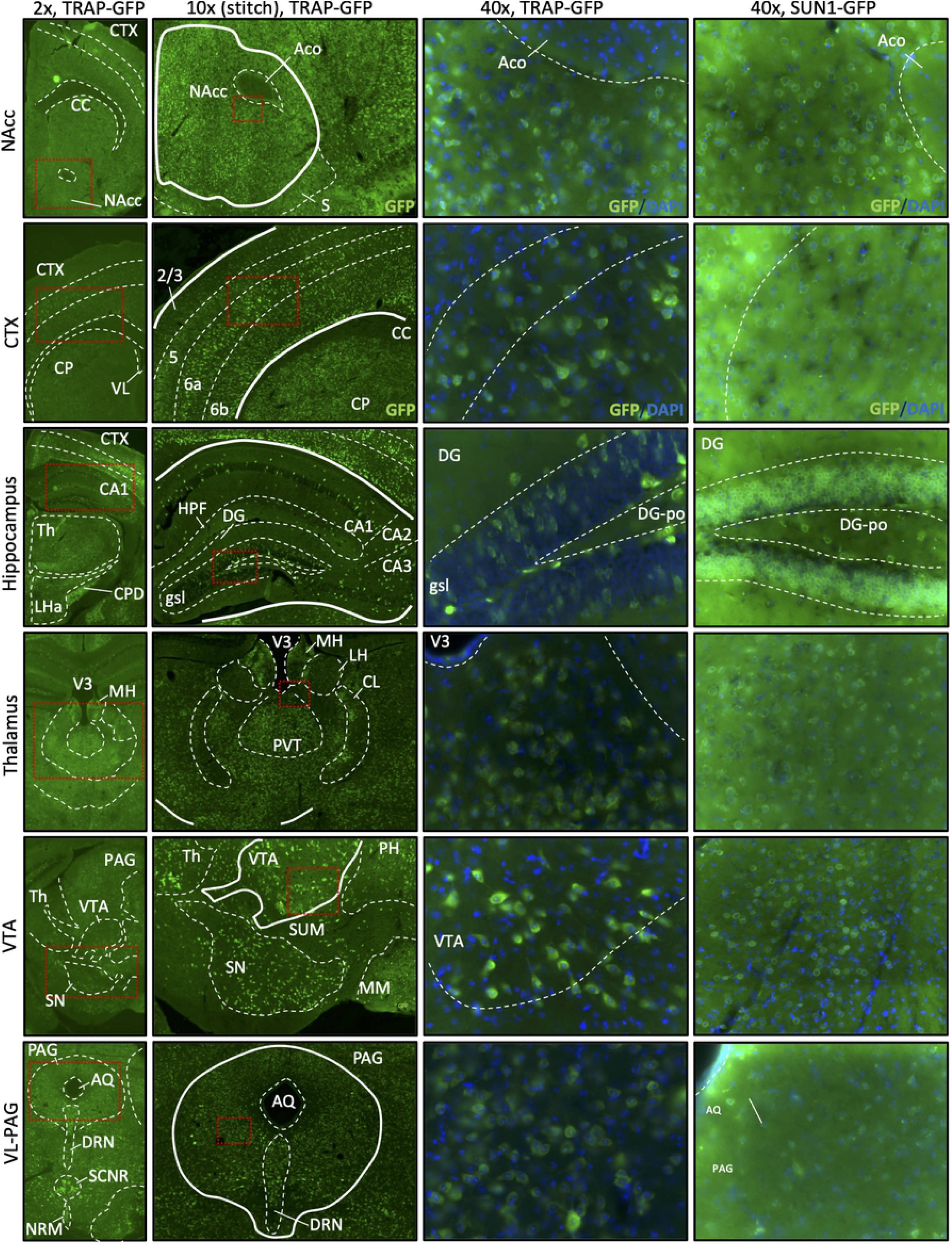
Imaging brain tissue from GFP-reporter mice crossed with the *Oprm1-Cre* line confirms functional Cre-mediated recombination. Recombinant GFP fluorescence reporting in Nucleus Accumbens (NAcc), Cortex (CTX), Hippocampus, Thalamus (Th), Ventral Tegmental Area (VTA) and ventrolateral periaqueductal-gray (VL-PAG). *Oprm1*-Cre x Rosa^LSL-GFP-L10a^ tissue (TRAP mice) and *Oprm1-Cre* x Rosa^LSLSun1-sfGFP^ (Sun1 mice) were imaged at 2x, 10x (z-stack/stitched image) and 40x magnification. Note that the ribosomal-GFP expression pattern is more dispersed and less defined in TRAP mice compared to the nuclear membrane staining in Sun1 mice. Slides were imaged using the BZ-X800 Viewer software in conjunction with a BZ-X Series automated Keyence microscope.

We further crossed the *Oprm1*^Cre/+^ line with the *Rosa*^LSLSun1-sfGFP^ reporter line, which expresses GFP at the inner nuclear membrane of Cre-expressing cells [18]. Upon imaging, brain regions expected to have high MOR density, similar to the results found with the Rosa^LSLSun1-sfGFP^ reporter, exhibited a high GFP signal (Fig 4). When visualized at 40x magnification, the distinct green ring around the DAPI-stained nuclei indicates proper Cre recombination and production of GFP from the *Oprm1*^Cre/+^;*Rosa*^LSLSun1-sfGFP^ line. Together, these imaging data demonstrate that *Oprm1*^Cre/+^ animals express functional Cre recombinase in MOR-cell types.

### Oprm1^Cre/+^; Rosa^LSL-GFP-L10a^ mice allow for the investigation of μOR cell-type specific changes in withdrawal

Actively translating mRNA from MOR-expressing cells was isolated from striatum of *Oprm1*^Cre/Cre^;*Rosa*^LSL-GFP-L10a^ mice undergoing spontaneous withdrawal following chronic morphine treatment of escalating doses (or saline control) and TRAP-sequencing data was assessed for cell-type specificity. Normalized gene counts were averaged across all TRAP-sequencing samples and the 10,000 most highly expressed TRAP-sequencing genes were analyzed for GO term enrichment using a reference data set of all possible genes (Fig 5A). The top GO terms were related to neural functions, such as synaptic transmission and organization, neuronal projections, and transport. To identify which cell types in the striatum were enriched in this TRAP-sequencing gene set, we performed hypergeometric tests to determine enrichment of cell-type specific gene lists from 3 separate published studies (Fig 5B-D). There was significant enrichment for all 3 neuron-specific gene list in our TRAP-sequencing data set (*p<0.05). Notably, there was very little enrichment for gene signatures from cell types other than neurons. When the same statistical tests were performed using publicly available bulk-sequencing data from the mouse striatum [19,20] there was significant enrichment for multiple non-neuronal cell types including microglia, astrocytes and oligodendrocytes (Fig 5B-D). Thus, as expected, the *Oprm1*^Cre/Cre^; *Rosa*^LSL-GFP-L10a^ model enriches for neuronal transcripts.

**Figure 5.**
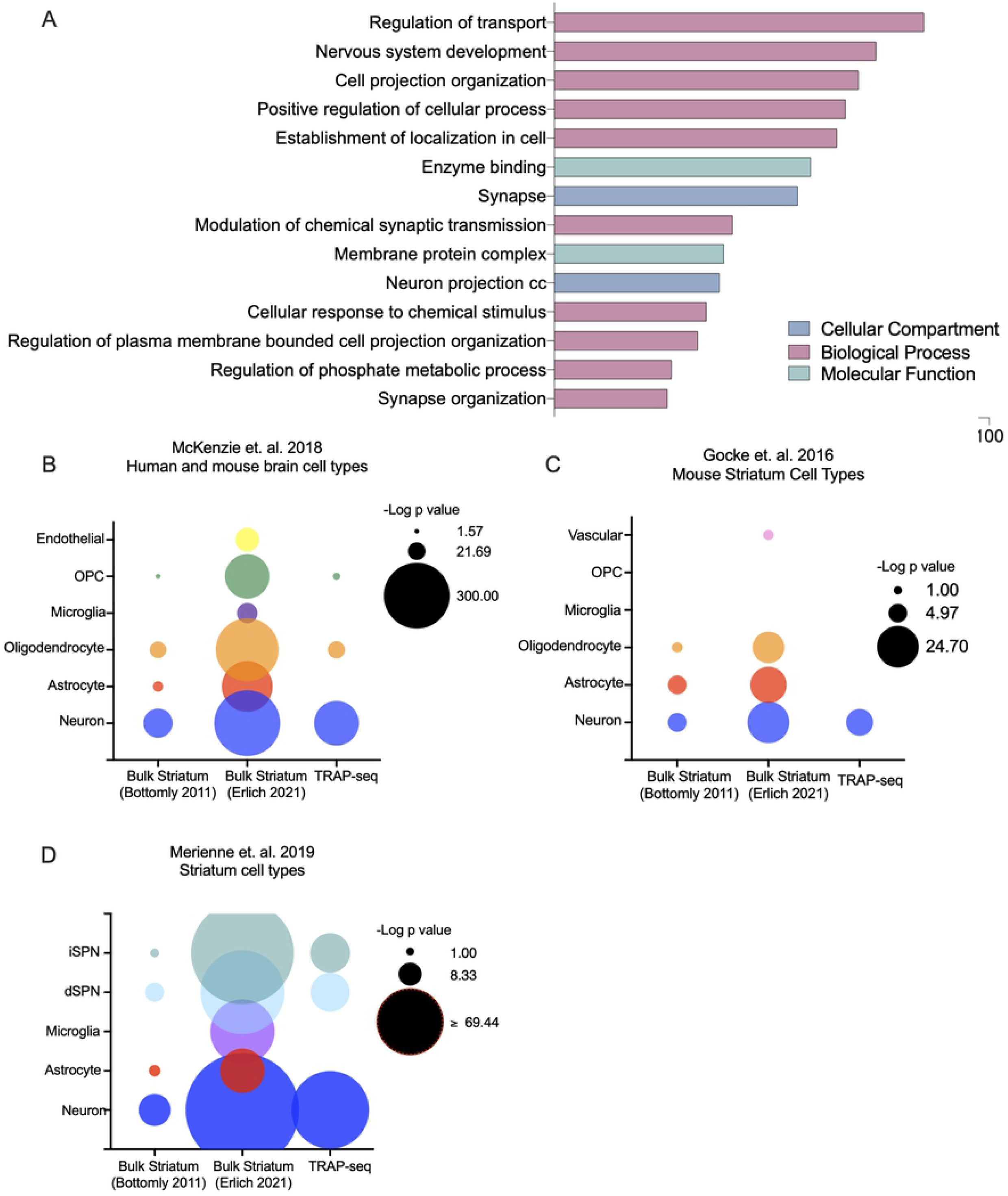
TRAP-sequencing in Oprm1-T2A-Cre striatum enriches for neuronal cell types. (A) GO analysis for enrichment in top 10,000 genes from TRAP-seq. The top 10,000 expressed genes (by average normalized gene count across all samples) were analyzed for GO term enrichment using goProfiler. All significant (p<0.05) GO terms were then reduced using Revigo. Top GO term categories are shown here. (B) Brain cell type enrichment. Enrichment for genes characteristic of brain cell types (McKenzie et. al. 2018) was calculated for using a hypergeometric test. Data sets from each sequencing experiment included the top 10,000 expressed genes (by average normalized gene count across control samples for bulk sequencing experiments, and across all samples for TRAP-sequencing experiment). (C and D). Striatum cell type enrichment. Enrichment for genes characteristic of striatum cell types was calculated in an identical manner for cell-specific gene lists from Gocke et. al. 2016 (C) as well as gene lists from Merienne et. al. 2019 (D).

TRAP-sequencing data from *Oprm1*-Cre mice undergoing withdrawal was compared to TRAP-sequencing data from saline-treated *Oprm1*-Cre controls (Fig 6). There were over 2,000 upregulated genes in the withdrawal group compared to the saline-treated controls (Fig 6A), while only a few, mostly uncharacterized genes were down-regulated. Therefore, we focused our subsequent analysis on upregulated genes. Upregulated genes included several previously implicated in withdrawal, such as *Plin4* [21]. Additional upregulated genes included *Enpp2, Atp1a2, Apoe* and *Slc1a3,* which play a role in psychiatric conditions often related to withdrawal, such as anxiety and depression [22–26]. Among the significantly upregulated genes, there was a significant enrichment of GO terms related to synapse and neuron function, including ion transport, enzyme binding, and neuronal development (Fig 6B).

**Figure 6.**
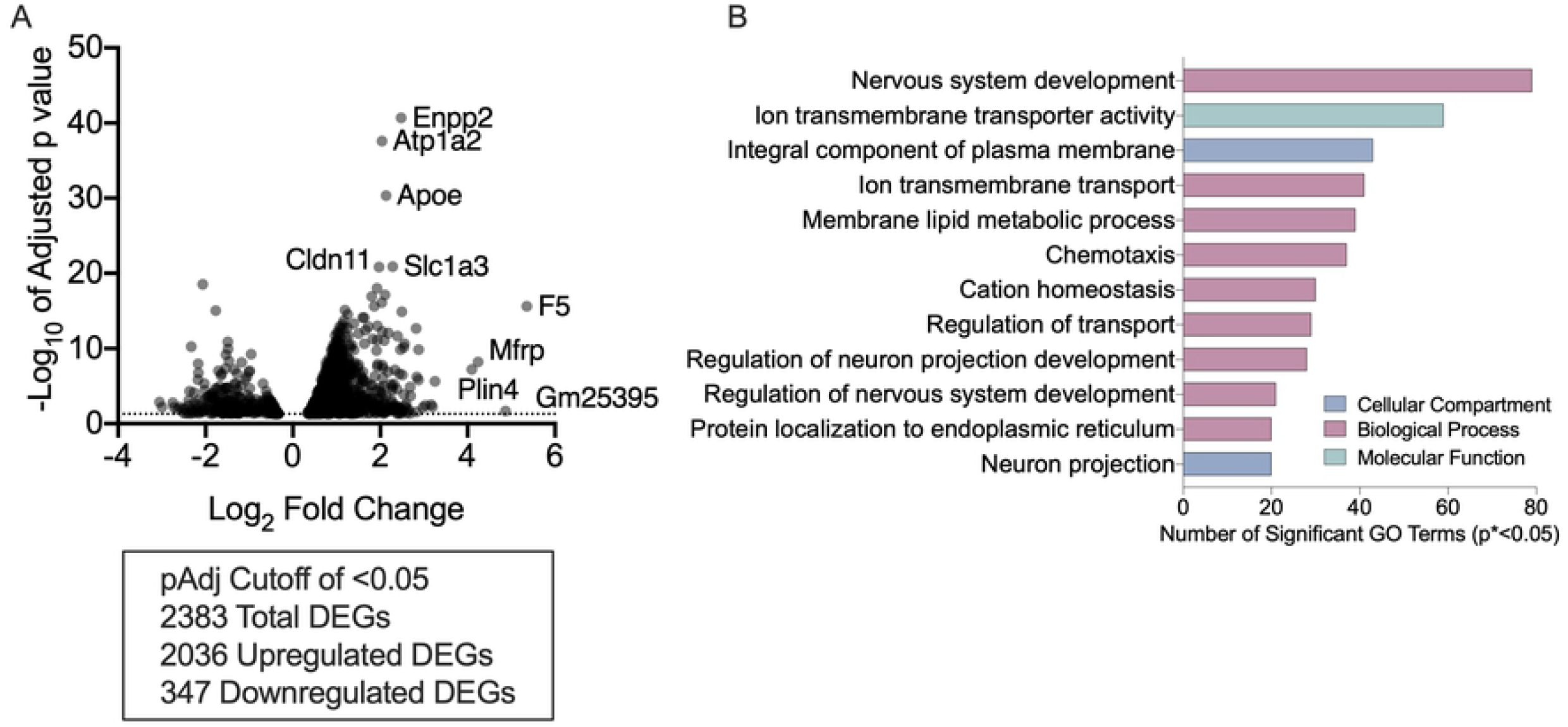
Chronic morphine withdrawal increases expression of membrane and synapse related genes in MOR+ cells in the striatum. (A) Differentially expressed genes in MOR+ cells in the striatum following chronic morphine withdrawal. (B) GO analysis for enrichment in DEGs (p<0.05) upregulated in MOR+ cells in the striatum following chronic morphine withdrawal. All significantly upregulated DEGs were analyzed for GO term enrichment using goProfiler. All significant (p<0.05) GO terms were then reduced using Revigo. Top GO term categories are shown here.

## Discussion

Here, we describe the derivation and characterization of a novel *Oprm1*-Cre mouse and demonstrate its utility as a tool to specifically interrogate MOR-expressing cells. Comparable levels of *Oprm1* expression and DAMGO binding across several different brain regions indicate no adverse effects of insertion of the T2A Cre construct into Exon 4 of the *Oprm1* locus. Additionally, this mouse line displayed no aberrant responses to acute or chronic morphine treatment. Together with binding data demonstrating similar density of MOR-expressing cells in relevant brain regions, as well as functional Cre recombinase activity, these data suggest that the *Oprm1*-Cre mouse has intact MOR activity and expression, and that the presence of the T2A cleavable peptide does not disrupt endogenous MOR function.

The new *Oprm1*-Cre mouse described here can be used in various Cre-recombinase systems to activate, deactivate, isolate, manipulate, sequence, image, and characterize MOR-expressing cells and cell-subpopulations. Importantly, these mice exhibit expected opiate-mediated behaviors and therefore can be used in a variety of assays to elucidate the role MOR’s play in these behaviors.

As our construct does not have fluorescent proteins fused to the Cre recombinase or the MOR itself, the *Oprm1*-Cre mouse line described here can be employed in experiments that require the use of a floxed-GFP reporter, and we demonstrated the utility of this *Oprm1-Cre* mouse line by crossing it with one such floxed GFP tool. Actively translating mRNA from the striatum of *Oprm1*^Cre/+^; *Rosa*^LSL-GFP-L10a^ mice was isolated via TRAP and sequenced. MOR expression in the striatum is enriched in clusters of medium spiny neurons (MSNs) that define the striosome compartment, however other cell types, such as astrocytes, oligodendrocytes and microglia, have been reported to express MORs as well [27,28], though the latter is controversial [29]. We found an enrichment of neuron and synapse-related GO terms and neuron-specific genes in our TRAP-sequencing data consistent with this. This contrasts with enrichment of several other brain cell types in bulk-sequencing experiments from mouse striatum. This suggests that the TRAP-sequencing with *Oprm1*-T2ACre did in fact isolate specifically neurons from the striatum.

Our striatal TRAP-sequencing data set provided insight into differentially expressed genes (DEG’s) in *Oprm1-expressing* cells between animals going through spontaneous morphine withdrawal and controls. Genes induced during withdrawal included several previously shown to play roles in substance abuse, affective disorders and morphine-induced changes in behavior and physiology. For example, the analgesic potency of morphine has been functionally attributed to *Atp1a2* (Na^+^, K^+^-ATPase α2 subunit) gene expression in the striatum, and the discovery of *Atp1a2* upregulation in our TRAP-sequencing data is of note considering that rodents under withdrawal experience hyperalgesia in the hot plate analgesia assay [30]. Additionally, mice with decreased *Atp1a2* activity display augmented fear/anxiety behaviors and enhanced neuronal activity [24]. Similarly, *ApoE,* activated in Oprm1 positive cells in the striatum during morphine withdrawal, has been implicated in several neuropsychiatric disorders including depression and schizophrenia [25], which can be co-morbid with substance use disorder [31]. *Slc1a3*, encoding a GABA transporter, has also been formally associated to be more expressed in males with Manic Depressive Disorder [26] and is considered a candidate gene for targeting anxiety disorders_[23], was also activated in Oprm1-positive striatal cells following morphine withdrawal.

Previous mouse striatal withdrawal sequencing studies found upregulation of *Plin4* and related genes, indicating that mechanistically, morphine withdrawal may have an effect on lipid packaging and production in the brain [21]. Finally, we found increased *Enpp2* expression, which is a phosphodiesterase that has been found to be upregulated in several inflammatory states [22]. This is consistent with the role of opioid withdrawal in altering inflammation, which is emerging as an important component of opioid biology [32]. However, out TRAP experiment identified several novel genes such as *F5*, which was first was first identified as an mRNA expressed by activated but not resting T-lymphocytes. The neuronal expression of *F5* mRNA correlates directly with the size of the neuronal perikarya, the length of the axonal projection, or the extent of dendritic arborization, suggesting dynamic alterations in neuronal plasticity following chronic opioid exposure and withdrawal.

Several GO terms associated with transport of molecules across the plasma membrane were upregulated in the striatal TRAP-sequencing data, consistent with altered neuronal function during withdrawal. The glutamine uptake transporter, *GLT1,* for example, has been shown to decrease expression in the NAc during short-term and long-term cocaine withdrawal, indicating that lower GLT1 expression may contribute to drug-seeking behavior post-withdrawal [33]. In terms of ion transport specifically, morphine acts in a concentration-dependent manner to reduce basal transmural potential differences and short-circuit currents in mouse jejunum *in vitro* [34].

K^+^-permeable channels in amygdalo-hippocampal neurons can decrease inhibition induced by morphine after chronic morphine treatment, and show firing activation post withdrawal [35]. We also found significant upregulation in GO terms associated with neuronal projections and nervous system development. Morphine withdrawal symptoms have previously been linked to neuronal differentiation and development, via the neurotrophin-3 (NT-3) neurotrophic factor [36], consistent with the TRAP mRNA sequencing data that revealed upregulation of these related GO-terms during morphine withdrawal. Additionally, in cell culture studies, morphine exposure has been shown to alter neuronal differentiation and neuronal projection directly [37,38].

Recently, several other Cre-driver strains of mice have been developed, which allow for cell type-specific Cre-recombinase expression in *Oprm1*-expresssing cells. A mouse that includes a T2A-cleaved, GFP-Cre recombinase fusion protein under control of the *Oprm1* promoter was used to ontogenetically activate *Oprm1*-expressing neurons in the VTA to induce avoidance behavior [39]. A second line has been generated with a tamoxifen-inducible *Oprm1-* driven Cre recombinase to enable temporospatial control of genetic manipulation of MOR expressing cells in the brain [40]. In a third line, Cre-GFP was inserted 5’ of the initiation codon in the first coding exon of the *Oprm1* gene, creating a null allele; these mice, when homozygote, no longer respond to certain endogenous or exogenous opioid ligands., making behavior assessments impossible. A fourth line was recently used to map molecular signatures of neuronal subdivisions within the striatum, based on their MOR expression [41]. In this case the *Oprm1*-Cre mouse was crossed with a Cre-dependent reporter line, Oprm1-Cre;H2B-GFP [42] allow identification of the RNA expression profile in single nuclei from MOR neurons.

Molecular characterization of neuron subtypes is becoming critical as technologies evolve to allow single-cell RNA sequencing, live cell imaging and cell-type specific behavior manipulation. Thus, having available tools that allows these technologies to use with opioid biology is timely. In addition, MOR-Cre lines can be used not only in combination with reporter mouse lines, but also to conditionally mutate a variety of genes specifically in *Oprm1-* expressing cells. This will allow for a finer examination of the functional role of genes regulated following opioid exposure and could provide insights on potential therapeutic targets.

## Acknowledgments

We wish to thank Drs. Catherine May, and Klaus Kaestner for CRISPER design protocols. We thank Dr. Jean Richa for embryo injections and Julia Noreck for genotyping assistance and colony maintenance.

